# Pathogens pull hardest in the coevolutionary arms-race to determine age-specific transmission biases

**DOI:** 10.1101/2025.11.19.689276

**Authors:** Samuel V. Hulse, Emily L. Bruns

## Abstract

Juveniles across a wide range of species are more susceptible to pathogenic infection than their adult conspecifics, often driving epidemic spread. While high juvenile susceptibility can be related to developmental timing, recent theoretical studies have found that hosts often evolve high juvenile susceptibility given a trade-off with other host traits. However, most research on age-specific infection has focused solely on host evolutionary dynamics, and the effect of pathogen evolution or coevolution on juvenile infection rates remains unknown. Here, we developed a coevolutionary adaptive dynamics model to investigate age-specific infectivity and contrast the outcome of coevolution with host evolution and pathogen evolution alone. We found strong differences in the evolutionary dynamics of hosts and pathogens in the absence of coevolution. While host evolution often favoured genotypes with intermediate levels of juvenile and adult resistance, pathogen evolution selected for extreme specialists. When hosts and pathogens were allowed to coevolve, the dynamics more closely resembled the pathogen dynamics; pathogens rapidly evolved to specialize on infecting a specific age class and this drove reciprocal specialization in host resistance. Our results illustrate that coevolution has the potential to shape age-specific patterns of infection that impact disease spread.

## Introduction

Juvenile hosts are typically more susceptible to pathogenic infection than adults (1–4). This age-specific susceptibility has important ecological consequences, and it has long been understood that age-structure has a strong impact on epidemiological outcomes (5–8). In natural populations, pathogen persistence can even depend on highly susceptible juveniles (9). The high susceptibility of juvenile hosts therefore has the potential to affect pathogen evolution: susceptible juveniles can offer pathogens a path of least resistance, allowing infectious agents to evade stronger aspects of adult immune defences. However, while several studies have examined the evolution of age-specific resistance in hosts, very few have investigated the evolutionary and coevolutionary dynamics of pathogen age-specific infectivity.

For hosts, juvenile susceptibility was long thought to be the byproduct of developmental constraints on immunity (10,11). However, recent theoretical and empirical work has demonstrated that juvenile susceptibility can be evolutionarily advantageous for the host, given a trade-off with host fecundity or mortality (4,12,13). However, all models of age-specific resistance evolution in the host have assumed a static, non-evolving pathogen (12,13), and likewise, age-specific pathogen evolution models have assumed a static host (14). Given the importance of coevolutionary dynamics on other aspects of host-pathogen persistence and prevalence (15–17), feedbacks between host and pathogen evolution may potentially upend previous assumptions about the evolution of juvenile and adult susceptibility.

Canonical host-pathogen coevolution models typically assume a single mechanism of resistance for the host, and a single corresponding mechanism of infectivity for the pathogen (15,17). However, different mechanisms often underly host resistance at juvenile and adult stages (1,4,18,19). For example, Bruns and Lesser found that a host’s resistance level at the juvenile stage was not predictive of their resistance level at the adult stage. In such cases, both hosts and pathogens must carefully balance investment in infectivity or defence between each pathway. Given multiple infection mechanisms, coevolution then becomes a two-front arms race, where both hosts and pathogens may face constraints in how much they can invest in either pathway. When these transmission pathways are associated with host age, coevolutionary dynamics may further depend on host life history or other physiological constraints associated with age.

While previous host evolution models show that selection broadly favours juvenile susceptibility and adult resistance, it is unclear whether this trend will hold in a coevolutionary context. Recently, Iritani et al. developed an evolutionary model where pathogens could alter their investment in both juvenile and adult infectivity, at the cost of increased virulence (14). Analogous to the trends found in host evolution models, their pathogen evolution model found that pathogen evolution broadly results in greater investment in juvenile infectivity. If selection favours the same outcomes in a coevolutionary context as in the non-coevolutionary context, both host and pathogen evolution will lead to higher transmission in juveniles. Alternatively, coevolution could alter selection on either the host or the pathogen, so that host and pathogen evolution counteract each other. In the first scenario, the combined effects of host and pathogen evolution will lead to extreme discrepancies in juvenile versus adult transmission, whereas in the second scenario, transmission will be more balanced across age classes.

To investigate the impact of host-pathogen coevolution on age-specific transmission dynamics, we developed an adaptive dynamics-based model. Because we wanted to directly compare host and pathogen investment in resistance and infectivity at different life stages, we assumed a trade-off between juvenile and adult resistance for the host, and between adult and juvenile infectivity for the pathogen. We first investigated how hosts and pathogens evolved in a non-coevolutionary context and then compared these outcomes to a full coevolutionary model. As earlier models have highlighted the importance of host life history in resistance evolution (Ashby & Bruns, 2018; Buckingham et al., 2023; Iritani et al., 2019), we test the effects of host maturation rate as well as the impact of hosts with intrinsically higher adult resistance.

## Methods

### (a) Model Description

We examined the evolutionary dynamics of a sterilizing disease with density-dependent transmission. In our model, we divide the host population into susceptible (*S*) and infected (*I*) compartments, both of which have a juvenile (*S*_*j*_, *I*_*j*_) and adult (*S*_*a*_, *I*_*a*_) stage. We assume that only susceptible adults are capable of reproduction (infection causes sterility but no additional mortality) and that transmission is density-dependent with no recovery. The transmission rate depends on the life stage of a susceptible host, as well as host resistance, and pathogen infectivity. We assume two life stages, juveniles and adults, where hosts transition from the juvenile to adult stage at rate *m*, referred to as the maturation rate. The dynamics of our model are loosely based on the sterilizing fungal pathogen “anther-smut” that infects plants in the carnation family. Host plants are significantly more susceptible at the juvenile stage than the adult stage (1), and resistance at this stage is known to be costly (4). Pathogen strains also vary in their ability to infect juvenile and adult hosts (Slowinski et al., *in review*), and importantly, host and pathogen generation times are equivalent. These dynamics are given by the following system of ordinary differential equations (Eqns. 1-4).

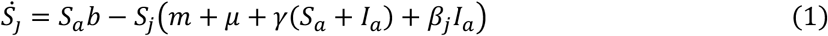

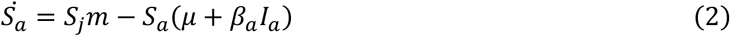

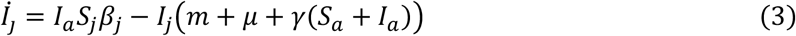

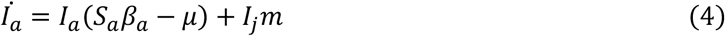

Host demographics are controlled by the birthrate, *b*, the maturation rate, *m*, the natural deathrate, μ (equal for all stages and infection statuses), and the coefficient of density-dependent regulation, γ (which acts only on juveniles and imposes a carrying capacity). Pathogen transmission is controlled by the age-specific transmission rates to juveniles, *β*_*j*_, and adults, *β*_*a*_. We further assume that transmission rates at each age are determined by both the host’s age-specific resistance (*h*_*j*_, *h*_*a*_), and the pathogen’s age-specific infectivity (*p*_*j*_, *p*_*a*_). The juvenile (*β*_*j*_) and adult transmission (*β*_*a*_) are therefore given by:

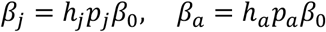

Here, *β*_0_ represents that baseline level of transmission, which is the same for juveniles and adults. Later, we investigate the effects of reducing the baseline transmission level for adults.

Our goal was to understand the outcome of different selection pressures shaping host and pathogen allocation to juvenile and adult resistance and infectivity, and how these changed under coevolution. We therefore assumed a reciprocal trade-off between juvenile and adult resistance in the host (*h*_*j*_, *h*_*a*_) and between juvenile and adult infectivity in the pathogen (*p*_*j*_, *p*_*a*_). In this way, the host’s resistance trade-off and the pathogen’s infectivity trade-off are equivalent, allowing us to contrast how selection induced by stage-specific transmission varies across hosts and pathogens. Because higher adult resistance is widely observed in nature, we investigated scenarios where adults have a higher baseline level of resistance than juveniles, given by *z*, the adult resistance advantage. Here, adults have a higher baseline resistance when *z* > 1. For simplicity, we did not include a similar modifier for the pathogen, as this would be equivalent to modifying *z*. In our simulations, we implement these trade-offs via a direct reciprocal trade-off, given by 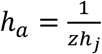 for the host, and 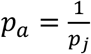 for the pathogen (we do not assume an a priori juvenile infectivity advantage for the pathogen). This trade-off function assumes that the While evidence in support of any particular trade-off function is scare, recent theoretical and empirical studies have found support for accelerating trade-off functions in costly traits (20–22).

**Figure 1.**
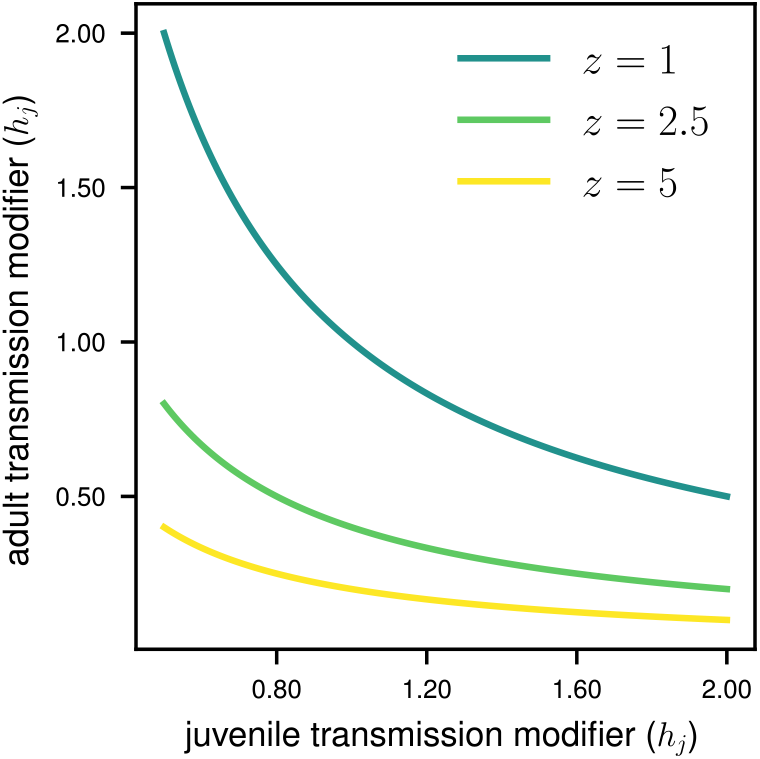
Trade-offs between adult and juvenile resistance. The trade-off function for pathogen infectivity is always *z* = 1, as we do not consider a separate pathogen infectivity bias parameter.

### (b) Evolutionary Invasion Analysis

We used the adaptive dynamics framework to implement host and pathogen coevolution. Here, we assume a haploid, asexual population. With adaptive dynamics, new mutants are iteratively introduced into a population with existing genotypes until an evolutionary equilibrium is reached. While we can derive analytical expressions for the fitness of mutant host and pathogen genotypes invading an established population (see Supplementary Information), this depends on the equilibrium prevalence of infected individuals, for which we could not find an analytical solution. We therefore used numerical simulations to investigate the evolutionary and coevolutionary outcomes of age-specific resistance and infectivity.

We began each simulation with a monomorphic host and pathogen population, where the initial genotypes had equal investment across age classes (*h*_*a*_ = *h*_*j*_, *p*_*a*_ = *p*_*j*_). Both hosts and pathogens were able to evolve up to a two-fold increase in adult or juvenile resistance or infectivity (*h*_*a*_, *h*_*j*_, *p*_*a*_, *p*_*j*_ ∈ [0.5, 2]), subject to the reciprocal trade-offs described earlier. We limited the range of evolvable resistance and infectivity values to prevent extreme transmission values for a single stage in the pathogen evolution model, and to prevent the host from effectively eliminating a transmission pathway in the host evolution model.

We began with a population of 100 susceptible adults, and 10 susceptible juveniles as well as 10 infected adults and 1 infected juvenile. We computed numerical solutions, from *t* = 0 to *t* = 5000 (this was deemed sufficient to approach equilibrium, as judged by visually analysing solutions across parameter space). At this point, we introduced mutants by shifting 5% of host and pathogen genotypes to mutant genotypes. For the host, these new mutants had slightly varied investments in adult versus juvenile resistance while mutant pathogens had slightly different adult and juvenile infectivity investments. As our trade-off function was non-linear, the magnitude of this difference increased as hosts and pathogens became more specialized. Specifically, values for resistance and infectivity were sampled from 2^*i*^, where *i* ranges from-1 to 1 in increments of 0.02. We then calculated new numerical solutions with the new mutants, to determine whether they had a higher fitness, or coexisted with the endemic genotypes. For each parameter set investigated, we then repeated this process for 200 iterations until both hosts and pathogens reached an evolutionary equilibrium.

We first tested how hosts allocate resources between juvenile and adult resistance given a non-evolving pathogen. Next, we considered the inverse case, where the pathogen can evolve different allocations between adult and juvenile infectivity given a non-evolving host. We then contrasted these outcomes with a coevolutionary model, in terms of both the evolutionary responses of hosts and pathogens, as well as the effects on pathogen transmission across age classes. For each of these evolutionary scenarios, we explored the role of maturation rate and the adult resistance advantage on evolutionary and coevolutionary outcomes of age-specific transmission. We tested maturation rates ranging from 0.25 to 1 and adult resistance advantage values ranging from 1 to 10 (we did not observe major qualitative changes in evolutionary outcomes beyond this range). For each set of parameters, we ran simulations with only host evolution, simulations with only pathogen evolution, and finally, coevolutionary simulations.

To investigate the impacts of host and pathogen evolution on host demographics and transmission, we calculated the stage-specific transmission ratio, based on the proportion of juveniles at evolutionary equilibrium (including infected individuals), the proportion of infected individuals (both adult and juvenile) and the degree of transmission. We defined the transmission ratio as 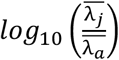, where 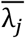 is the juvenile force of infection and 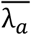 is the adult force of infection. With this metric, negative values indicate more transmission to adults and positive values mean more transmission to juveniles. The force of infection is calculated by the following:

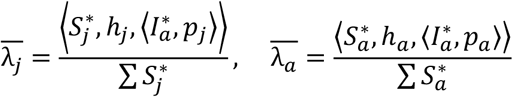

Here, 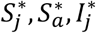 and 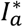 represent the equilibrium vectors of host and pathogen genotype abundances.

## Results

### (a) Host Evolution: Mathematical Analysis

The fitness of a new mutant host genotype can be determined using the next generation matrix method (23), yielding the following expression (see Supplementary Information for derivation) for the fitness of a mutant host invading a population at equilibrium:

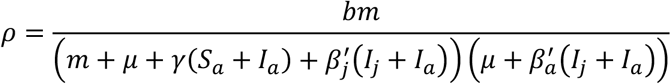

A mutant host genotype with transmission rates 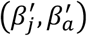 can invade if *ρ* > 1. For simplicity, we write this equation in terms of transmission, not resistance. Assuming equal pathogen investment in infectivity (*p*_*a*_ = *p*_*j*_), the relationship between transmission and resistance is 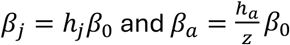. Due to this reciprocal relationship between *h*_*a*_ and *h*_*j*_, terms where adult and juvenile transmission terms are multiplied are proportional to 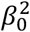, and thus have a small impact on the overall invasion fitness. As 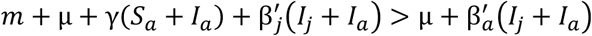 when β_*j*_ = β_*a*_, a mutant with a proportional increase in investment in adult resistance will have a greater invasion fitness than a mutant with higher investment in juvenile resistance. As we assume nonlinear trade-offs between adult and juvenile resistance, this does not necessarily lead to directional selection as at some point the benefits of further decreases in 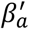 will be offset by the corresponding increase in 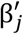;

### (b) Pathogen Evolution: Mathematical Analysis

As with the invasion fitness of a mutant host genotype, we used the next generation matrix method to determine the fitness of a mutant pathogen invading a population at equilibrium (see Supplementary Information for derivation):

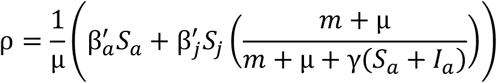

Assuming equal host investment in resistance (*h*_*a*_ = *h*_*j*_), the relationship between transmission and infectivity is 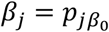 and 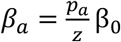. Unlike with host resistance evolution, where the transmission terms interact multiplicatively, with pathogen evolution, the transmission terms are additive. Since 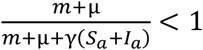, a proportional increase in 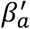 will have a larger impact on invasion fitness that 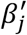. Therefore, either *z* must be large, or the susceptible host population must have a high proportion of juveniles for selection to favor juvenile infectivity.

### (c) Numerical analysis

Our evolutionary invasion analysis revealed markedly different evolutionary dynamics between hosts and pathogens. We first describe the dynamics for a pathogen when the host has a moderately fast maturation rate (*m* = 0.75) and higher relative adult resistance (*z* = 5), as has been observed in natural populations of wild carnations (9). We found that selection in the model with only host evolution, a moderate increase in adult resistance was generally favoured (Fig. 2A). However, in the model with only pathogen evolution, selection favoured full specialization on adults (Fig. 2B). In the coevolutionary model, the pathogen remained under strong directional selection for adult resistance, which led the host to fully specialize in adult resistance to counter pathogen evolution (Fig. 2C). Ultimately, this meant that in our coevolutionary model, the transmission ratio between adults and juveniles did not change greatly from its initial conditions, as host and pathogen evolution counteracted each other.

**Figure 2.**
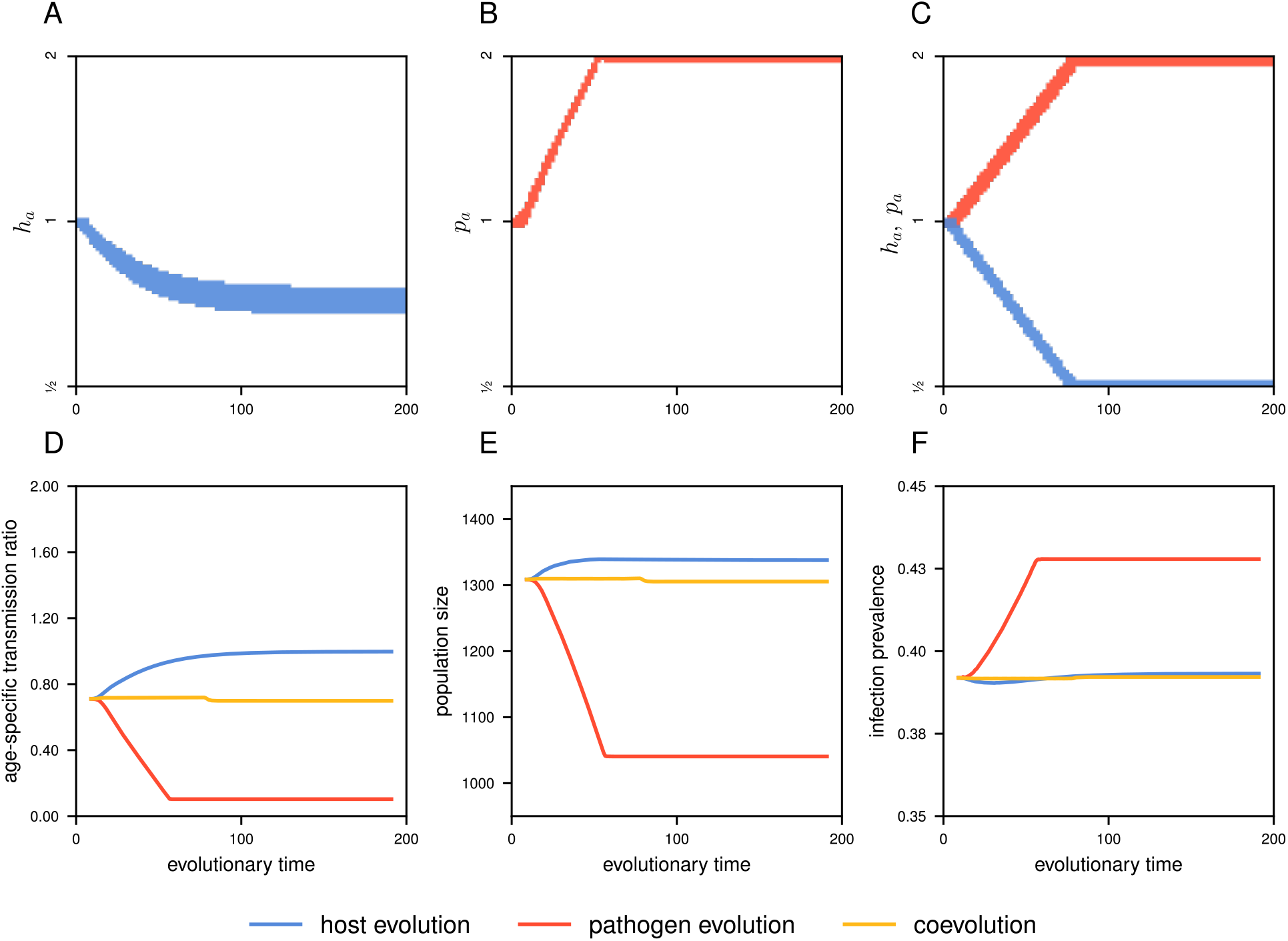
Example evolutionary outcomes with a moderately fast maturation rate (*m* = 0.75) and intermediate adult resistance advantage (*z* = 5). Results for (A) host evolution mode, (B) pathogen evolution model and (C), coevolutionary model. For A-C, the blue lines represent the host’s resistance modifier (*h*_*a*_), where values greater than 1 represent increased investment in adult resistance, and the red lines represent the pathogen’s infectivity modifier (*p*_*a*_), where values greater than 1 represent increased investment in adult infectivity. (D): The host’s juvenile transmission bias where values below zero indicate a greater transmission to adults than juveniles. (E): The total population density, including both age classes and infection statuses. (F): The proportion of all hosts that are infected. Parameters: *β*_0_ = 0.001, *γ* = 0.01, *b* = 1, *μ* = 0.2.

The different evolutionary outcomes in turn affected transmission dynamics, host age structure, and the overall levels of disease prevalence (Fig. 2D-F). When only hosts evolved, they invested in adult resistance, which shifted the burden of infection to juveniles (Fig. 2D, blue line). This led to a modest increase in the overall population density (Fig. 2E) but had little effect on the infection prevalence (Fig. 2F). Pathogen evolution (with no host evolution) had the opposite effect: a shift in transmission towards adults (Fig. 2D, red line), a significant decrease in the total population denisty (Fig. 2E), and an increase in the infection prevalence (Fig. 2F). In the pathogen evolution model, the pathogen was able to fully specialize on either adult or juvenile infectivity, leading to larger impacts on host demographics and the infection prevalence than in either the host evolution or coevolutionary models. The coevolutionary model generally led to outcomes between the host evolution and pathogen evolution models in terms of the transmission (Fig. 2D) bias and the population density (Fig. 2E). However, the total infection prevalence did not differ substantially between the host and coevolutionary model (Fig. 2F). This was due to the higher population density in the host evolution model, with increased the density-dependent transmission pressure.

### (d) Effect of the adult-juvenile resistance ratio and maturation rate

To determine the generality of our results to variation in maturation rates, which impact host age structure, and the adult resistance advantage, we ran simulations across a broad range of *m* and *z* values (Fig. 3). When only hosts were allowed to evolve, selection generally favoured investment in adult resistance (at the cost of increased juvenile susceptibility; blue shaded region: Fig. 3A). Both fast maturation rates and low adult resistance advantage increased selection for adult resistance, with a smooth transition in the equilibrium genotype. Shorter juvenile periods (higher values of *m*) decreased the likelihood of infection during this period, thus leading to increased investment in adult resistance over juvenile resistance. When adults were inherently more resistant (higher *z* values), the marginal benefit of increased adult resistance declined, thus selection favored higher juvenile resistance.

**Figure 3.**
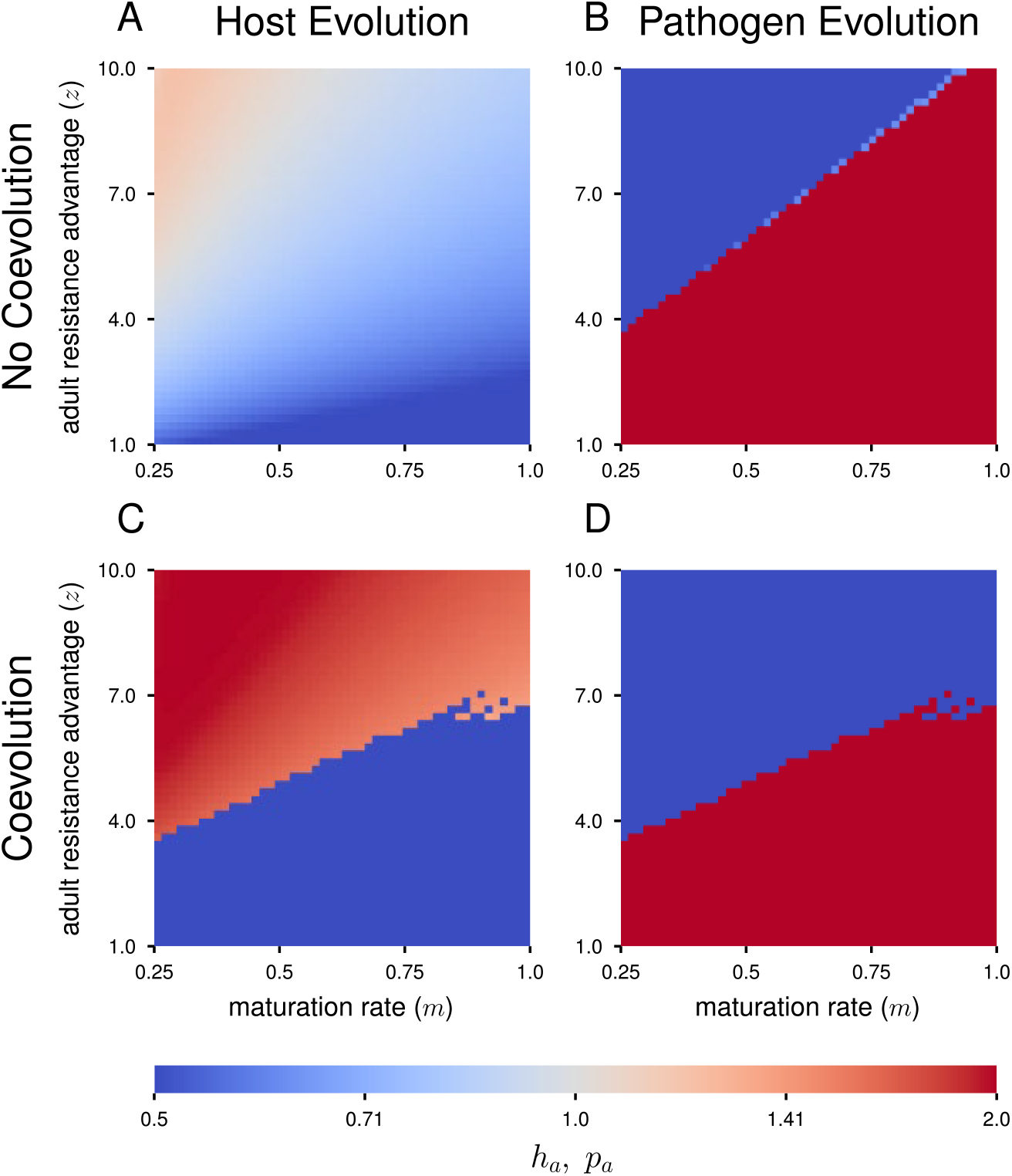
Evolutionary dynamics for the host evolution model (A), pathogen evolution model (B) and coevolutionary models (C-D). Blue hues indicate hosts investing in adult resistance (A, C) or pathogens investing in juvenile infectivity (B, D) (leading to high juvenile transmission) while red hues indicate hosts investing in juvenile resistance or pathogens investing in adult infectivity (leading to high adult transmission). Other parameters: *β*_0_ = 0.001, *γ* = 0.01, *b* = 1, *μ* = 0.2.

When only pathogens could evolve, we found two distinct regions of parameter space where either full adult infectivity specialization evolved (blue region, Fig. 3B), or full juvenile specialization (red region, Fig. 3B). However, we found a small region where evolution led to a polymorphic pathogen population composed of both full adult specialists and full juvenile specialists (light blue region in between juvenile and adult specialist regions, Fig. 3B). The transitions between adult specialization, polymorphism, and juvenile specialization occurred abruptly, across a very small range of parameters, both in terms of the maturation rate and the adult resistance advantage. No parameters led to intermediate pathogen genotypes at equilibrium. Longer juvenile periods (lower values of *m*) and higher adult resistance advantage (*z*) led to increased investment in juvenile infectivity over adult infectivity.

Coevolution markedly changed the host evolutionary dynamics (Fig. 3C) Whereas in our host evolution model we saw a gradual shift from investment in adult versus juvenile resistance, in our coevolutionary model this shift is abrupt, with fewer parameters leading to intermediate genotypes. The transition in host evolution corresponds to regions where the pathogen transitions from specializing on adult infectivity to juvenile infectivity. Coevolution shifted the transition zone between juvenile and adult specialization for the pathogen, expanding the region that favours full juvenile specialization (blue region: Fig. 3D).

Without a strong adult resistance advantage (*z* = 1), the host maturation rate had little impact on transmission dynamics (Fig. 4A). Here, the host maximizes adult resistance, while the pathogen maximizes adult infectivity. However, with a higher adult resistance advantage (*z* = 5), the pathogen’s evolutionary strategy depends on the host maturation rate, with significant implications for transmission dynamics (Fig. 4B). As the maturation rate crossed the threshold for the pathogen to shift from juvenile specialization to adult specialization, significantly more transmission shifted to adults. This dynamic was also apparent in our coevolutionary model to a lesser extent, although reciprocal host evolution greatly reduced its effect (yellow line, Fig. 4B). Increasing the maturation rate also significantly increased the host population density (Fig. 4C-D). By shortening the juvenile period, hosts escaped the density-dependent regulation of juveniles more rapidly and were able to reproduce sooner. As pathogen transmission is density-dependent in our model, as the host population density increased, so too did the force of infection. As a result, the number of infected hosts (dashed lines, Fig. 4C-D) often increased faster than the overall population density (solid lines, Fig. 4C-D) as the maturation rate increased. While this effect trend occurred for both high and low values of adult resistance advantage, the overall infection prevalence was significantly reduced for higher levels of *z*.

**Figure 4.**
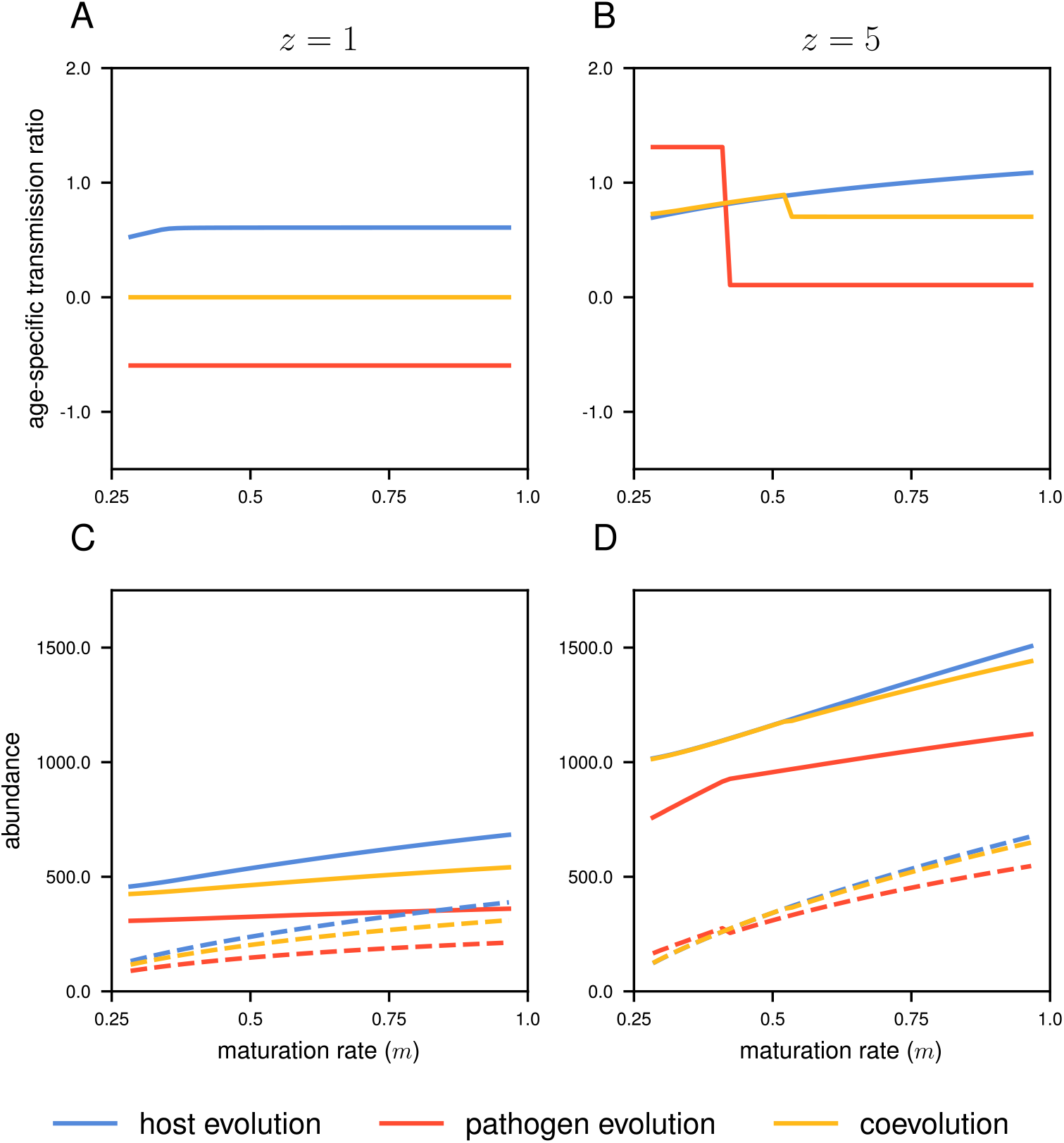
Effects of host maturation rate on transmission by juveniles and adults (A, B) and host abundance (C, D) for two different values of *z*, one where the pathogen maintains adult specialization (*z* = 1: A, C) and one where the pathogen shifts from juvenile to adult specialization (*z* = 5: B, D). Positive age-specific transmission ratios indicate more transmission to juveniles when negative values indicate more transmission to adults. For the host abundance (C, D), the solid lines represent the total population density, while the dashed lines represent the infected population density. Parameters: *β*_0_ = 0.001, *γ* = 0.01, *b* = 1, *μ* = 0.2.

## Discussion

We hypothesized that age-specific coevolution could result in two different outcomes: either direct antagonistic evolution at the same stage (increased adult resistance for the host and increased adult infectivity for the pathogen), or alternatively, the path of least resistance (increased adult resistance leads to increased juvenile infectivity). A common thread across all our models with pathogen evolution, and across all parameters was that the pathogen always evolves full specialization on either juveniles or adults. We found the pathogen’s “all-in” strategy induced a proportional response via host evolution, where the host evolved full specialization to counter the full specialization of the pathogen. Therefore, in our coevolutionary model, host and pathogen evolution acted antagonistically, where the effects of host evolution largely nullified the effects of pathogen evolution, and vice versa. In most cases, this led to more transmission occurring at the juvenile age, consistent with previous models of age-specific host resistance evolution (12,13). However, the magnitude of this effect is much less than would be the case if coevolution reflected combined outcomes of both the host evolution and pathogen evolution models.

The coevolutionary dynamics of our model resulted from fundamental constraints on the host: while the pathogen can persist with only one transmission pathway, host persistence depends on both juveniles and adults. When conditions favour one infection pathway over another, selection on the pathogen leads to an all-in strategy, while selection on the host leads to a more balanced approach. This dynamic is clear in our model: in both our pathogen evolution model and our coevolutionary model, we found that the pathogen was under strong directional selection for specialization. As the direction of selection is proportional to the ratio of juveniles to adults, without strong demographic feedback from infection, selection on the pathogen remains directional. While we did find demographic impacts on the host from pathogen evolution, these were never sufficient to induce balancing selection on the pathogen. While host evolution may generally lead to highly susceptible juveniles, host-pathogen coevolution may temper these trends via antagonistic coevolutionary forces. Given the sharp threshold across which pathogen evolution shifts specialization, shifting host life-histories may have small impacts until they suddenly have large impacts.

Our model assumes a direct trade-off between adult and juvenile resistance or infectivity. While few studies have directly tested this assumption, there is abundant evidence to suggest that adult and juvenile resistance can evolve independently (1,4). In this context, our approach can illustrate the relative strength of selection for resistance or infectivity at each stage, even if such biological trade-offs involve other factors, such as maturation rates or mortality. Furthermore, work in anther-smut fungal pathogens suggests that individual strains may face a trade-off between adult and juvenile infectivity on some host backgrounds (Slowinski et al., in review). Regardless of the precise trade-off, our model still demonstrates that the relative strength of selection for pathogen infectivity between adults and juveniles can shift far quicker than for host resistance. Pathogens can therefore respond more nimbly in how they allocate resources between multiple infection pathways. Importantly, this more rapid evolutionary response for pathogens occurred even though generation times were the same for hosts and pathogens.

In our model, coevolution allowed the host to regulate the effects of pathogen evolution, but in natural systems, the relative rate of evolution and standing genetic variation will determine whether directional selection on the pathogen can be arrested (24). While it is often assumed that pathogens can evolve faster than their hosts, owing to their shorter generation times, this is not always the case. For example, for a pathogen that is relatively static, such as highly selfing fungi (25,26), host evolution against a static pathogen may be a valid assumption. Conversely, for a more rapidly evolving pathogen such as a RNA virus, rapid pathogen evolution might lead to more pronounced demographic impacts on the host (27). These patterns may be more important for pathogens that have recently undergone host shifts, and both host and pathogen are far from evolutionary equilibrium.

While it has long been appreciated that the age-structure plays a critical role in the spread of disease, until recently this has received less attention from an evolutionary perspective. Whenever hosts rely on separate forms of resistance at juvenile and adult stages, pathogens also wage a multi-front coevolutionary arms race. Our results show that this arms race is more of a tug-of-war, where the pathogen determines the direction of the first pull.

## Ethics

This work did not require any ethical approval from a human subject or animal welfare committee.

## Data accessibility

All scripts used to generate model outputs are available as an electronic supplementary material.

## Declaration of AI use

We have not used AI-assisted technologies in creating this article.

## Authors’ contributions

S.V.H: conceptualization, formal analysis, software, visualization, writing – original draft, writing – review and editing. E.L.B.: funding acquisition, conceptualization, supervision, writing – review and editing.

## Acknowledgements

We would like to thank Janis Antonovics for his encouragement, feedback and many helpful discussions.

