## Supplementary Materials for "Pathogens pull hardest in the coevolutionary arms-race to determine age-specific transmission biases"

### 1 The Model

We model host infection with a compartmental model consisting of adults and juveniles for both the susceptible and infected classes. Only susceptible adults are capable of reproduction while only infected adults are capable of disease transmission. We define the dynamics of our model by the following system of ordinary differential equations:

$$\dot{S}_j = bS_a - (m + \mu + \gamma(S_a + I_a) + \beta_j(I_j + I_a))\, S_j \tag{1}$$
$$\dot{S}_a = mS_j - (\mu + \beta_a(I_j + I_a))\, S_a \tag{2}$$
$$\dot{I}_j = (I_j + I_a)\beta_j S_j - (m + \mu + \gamma(S_a + I_a))\, I_j \tag{3}$$
$$\dot{I}_a = (I_j + I_a)\beta_a S_a + mI_j - \mu I_a \tag{4}$$

Here, we assume that disease transmission is density-dependent, resulting in sterility without additional mortality. Host demographics are determined by $b$, the birth-rate, $\mu$, the death-rate, and by $\gamma$, the coefficient of density-dependent growth. Transmission to susceptible juveniles is determined by $\beta_j$ while transmission to susceptible adults is given by $\beta_a$.

#### 1.1 Transmission

##### 1.1.1 Infectivity and Resistance

We assume that disease transmission is a property of both the host (resistance) and pathogen (infectivity). We assume further that both of these factors depend on the age of the host. With this framework, the transmission coefficients are determined by

$$\beta_j = h_j p_j \beta_0$$
$$\beta_a = h_a p_a \beta_0$$

Where $h$ is the host resistance for either juveniles or adults and $p$ is the infectivity of the pathogen against either juveniles or adults. We assume that the baseline level of transmission $\beta_0$ is the same for both age classes.

##### 1.1.2 Trade-offs

We assume that host resistance at the juveniles stage trades-off with resistance at the adult stage. Likewise, pathogen infectivity against hosts at the juvenile stage trades-off against infectivity against adults. We assume a simple reciprocal relationship between adult and juvenile resistance, given by

$$h_a = \frac{1}{zh_j}, \quad p_a = \frac{1}{p_j}$$

This trade-off function has the property

$$f''(h_j) = \frac{2}{zh_j^3} > 0 \quad \forall h_j > 0$$

Thus, this is an accelerating, or convex trade-off function. We can then write adult transmission as

$$\beta_a = \frac{\beta_0}{zh_j p_j}$$

Where the parameter $z$ controls the shape of the trade-off for host resistance. If $z = 1$, then hosts can invest in either juvenile or adult resistance equally effectively, while if $z > 1$, host resistance is biased towards adults. This reflects that in many systems, adult have higher baseline levels of resistance against pathogens.

##### 1.2 Disease-Free Equilibrium

The disease-free equilibrium ($I_j = I_a = 0$) is given by the following.

$$\begin{aligned}
S_j^\star &= \frac{mb - m\mu - \mu^2}{m\gamma} \\
S_a^\star &= \frac{mb - m\mu - \mu^2}{\mu\gamma}
\end{aligned}$$

This also gives us the relation

$$S_j^\star = S_a^\star \frac{\mu}{m}$$

##### 1.3 Interior Equilibrium

We begin with the system at equilibrium:

$$\begin{aligned}
bS_a &= (m + \mu + \gamma(S_a + I_a) + \beta_j(I_j + I_a))\, S_j && (5) \\
mS_j &= (\mu + \beta_a(I_j + I_a))\, S_a && (6) \\
(I_j + I_a)\beta_j S_j &= (m + \mu + \gamma(S_a + I_a))\, I_j && (7) \\
(I_j + I_a)\beta_a S_a &= \mu I_a - m I_j && (8)
\end{aligned}$$

Using (2) and (4), we obtain

$$\begin{aligned}
& m(S_j - I_j) = \mu(S_a + I_a) \\
\implies \quad & S_a + I_a = \frac{m}{\mu}(S_j - I_j)
\end{aligned}$$

While using (1) and (3), we obtain

$$\begin{aligned}
bS_a &= (m + \mu + \gamma(S_a + I_a) + \beta_j(I_j + I_a))\, S_j \\
&= (m + \mu + \gamma(S_a + I_a))S_j + (m + \mu + \gamma(S_a + I_a))I_j \\
&= (m + \mu + \gamma(S_a + I_a))(S_j + I_j)
\end{aligned}$$

Combining these statements, this can be simplified to

$$\frac{bS_a}{S_j + I_j} = \frac{m\mu + \mu^2 + m\gamma}{\mu}(S_j - I_j)$$

However, we are unable to find an exact formula for the interior equilibrium.

##### 1.4 Stability of the Disease-Free Equilibrium

To evaluate the stability of the disease-free equilibrium, we take the Jacobian submatrix representing the host dynamics at equilibrium.

$$Df_h(x^\star) = \begin{bmatrix} -m - \mu - \gamma S_a & b - S_j\gamma \\ m & \mu \end{bmatrix}$$

This has eigenvalues

$$\lambda_1 = -\frac{bm + \mu^2 + \sqrt{b^2m^2 + 2bm\mu^2 + 4m\mu^3 + 5\mu^4}}{2\mu}$$
$$\lambda_2 = \frac{-bm + \mu^2 + \sqrt{b^2m^2 + 2bm\mu^2 + 4m\mu^3 + 5\mu^4}}{2\mu}$$

Clearly $\lambda_1 < 0$ for all positive parameter values, thus the stability only depends on $\lambda_2$. Therefore, the disease-free equilibrium is stable given $\lambda_2 > 0$ and $\mathcal{R}_0 < 1$.

#### 2 Host Evolutionary Dynamics

To examine the invasion potential of a mutant host genotypes, we append a mutant juvenile and adult compartment onto our system, and consider the subsystem composed of these compartments. Given a mutant genotype introduced at low frequency, this becomes

$$\dot{S}_j' = bS_a' - \left(m + \mu + \gamma(S_a + I_a) + \beta_j'(I_j + I_a)\right) S_j'$$
$$\dot{S}_a' = mS_j' - (\mu + \beta_a'(I_j + I_a))\, S_a'$$

The Jacobian submatrix is thus

$$Df(x) = \begin{bmatrix} -m - \mu - \gamma(S_a + I_a) - \beta_j'(I_j + I_a) & b - \gamma S_j' \\ m & -\mu - \beta_a'(I_j + I_a) \end{bmatrix}$$

##### 2.1 Invasion Fitness

Next, using the next-generation method, we have

$$J(x) = F - V = \begin{bmatrix} 0 & b \\ 0 & 0 \end{bmatrix} - \begin{bmatrix} m + \mu + \gamma(S_a + I_a) + \beta_j'(I_j + I_a) & \gamma S_j' \\ -m & \mu + \beta_a'(I_j + I_a) \end{bmatrix}$$

Therefore,

$$\begin{aligned} FV^{-1} &= \begin{bmatrix} 0 & b \\ 0 & 0 \end{bmatrix} \begin{bmatrix} \mu - \beta_a'(I_j + I_a) & -\gamma S_j' \\ m & m + \gamma(S_a + I_a) + \mu + \beta_a'(I_j + I_a) \end{bmatrix} \\ &= \begin{bmatrix} bm & b(m + \gamma(S_a + I_a) + \mu + \beta_a'(I_j + I_a)) \\ 0 & 0 \end{bmatrix} \end{aligned}$$

Which gives us

$$\rho(FV^{-1}) = \frac{bm}{\left(m + \mu + \gamma(S_a + I_a) + \beta_j'(I_j + I_a)\right)\left(\mu + \beta_a'(I_j + I_a)\right)}$$

Since $\rho(FV^{-1}) = 1$ when $\beta_j' = \beta_j$, we can use (1) to obtain

$$1 = \frac{bm}{b\frac{S_a}{S_j}(\mu + \beta_a'(I_j + I_a)} \Rightarrow m\frac{S_j}{S_a} = \mu + \beta_a'(I_j + I_a)$$

Substituting this back into our original expression, we get

$$\rho(FV^{-1}) = \frac{bS_a}{\left(m + \mu + \gamma(S_a + I_a) + \beta_j'(I_j + I_a)\right) S_j}$$

#### 2.2 Fitness Gradient

$$\omega(h_j) = \frac{bm}{(m + \mu + \gamma(S_a + I_a) + \beta_0 h_j(I_j + I_a))\left(\mu + \frac{\beta_0}{zh_j}(I_j + I_a)\right)}$$

$$\frac{\partial \omega}{\partial h_j} = \frac{\beta_0^2 mz(I_j + I_a)\left(\gamma(S_a + I_a) + m - \mu z h_j^2 + \mu\right)}{(I_a + \beta_0 I_j + \mu z h_j)^2\left(S_a(\beta_0 h_j + \gamma) + \beta_0 h_j S_j + \gamma S_a + m + \mu\right)^2}$$

As we are principally interested in the sign on the fitness gradient, we can reduce this to

$$\frac{\partial \omega}{\partial h_j} = \left(\gamma(S_a + I_a) + m + \mu - \mu z h_j^2\right)$$

### 3 Pathogen Evolutionary Dynamics

We begin by appending mutational compartments onto our model

$$\begin{aligned} \dot{I}_j' &= (I_j' + I_a')\beta_j' S_j - (m + \mu + \gamma(S_a + I_a))\, I_j' \\ \dot{I}_a' &= (I_j' + I_a')\beta_a' S_a + mI_j' - \mu I_a' \end{aligned}$$

The Jacobian of the mutant submatrix is then given by

$$Df(x) = \begin{bmatrix} \beta_j' S_j - m - \mu - \gamma(S_a + I_a) & \beta_j' S_j \\ \beta_a' S_a + m & \beta_a' S_a - \mu \end{bmatrix}$$

##### 3.1 Invasion Fitness

Using the next-generation method, we have

$$J(x) = F - V = \begin{bmatrix} \beta_j' S_j & \beta_j' S_j \\ \beta_a' S_a & \beta_a' S_a \end{bmatrix} - \begin{bmatrix} m + \mu + \gamma(S_a + I_a) & 0 \\ -m & \mu \end{bmatrix}$$

Therefore,

$$\begin{aligned} FV^{-1} &= \begin{bmatrix} \beta_j' S_j & \beta_j' S_j \\ \beta_a' S_a & \beta_a' S_a \end{bmatrix} \begin{bmatrix} \frac{1}{m+\mu+\gamma(S_a+I_a)} & 0 \\ \frac{m}{\mu(m+\mu+\gamma(S_a+I_a))} & \frac{1}{\mu} \end{bmatrix} \\ &= \frac{1}{\mu} \begin{bmatrix} \frac{(m+\mu)(\beta_j' S_j)}{m+\mu+\gamma(S_a+I_a)} & \beta_j' S_j \\ \frac{(m+\mu)(\beta_a' S_a)}{m+\mu+\gamma(S_a+I_a)} & \beta_a' S_a \end{bmatrix} \end{aligned}$$

Which gives us

$$\rho(FV^{-1}) = \frac{1}{\mu} \left( \beta_a' S_a + \beta_j' S_j \left( \frac{m+\mu}{m+\mu+\gamma(S_a+I_a)} \right) \right)$$

The invasion condition is thus

$$\mu < \frac{m+\mu}{m+\mu+\gamma(S_a+I_a)} \beta_j' S_j + \beta_a' S_a$$

A sufficient condition for invasion is thus

$$\mu < \beta_j' S_j + \beta_a' S_a$$

Using the tradeoff assumption, this becomes

$$\frac{\mu}{\beta_0} < p_j S_j + \frac{1}{p_j z} S_a$$

##### 3.2 Fitness Gradient

$$\frac{\partial \omega}{\partial p_j} = \frac{\beta_0}{\mu} \left( \left( \frac{h_j(m+\mu)}{m+\mu+\gamma(S_a+I_a)} \right) S_j - \frac{1}{z h_j p_a^2} S_j \right)$$
